# CUB domain containing protein 1 (CDPC1) is a target for radioligand therapy in castration resistant prostate cancer

**DOI:** 10.1101/2021.06.17.448569

**Authors:** Ning Zhao, Kai Trepka, Yung-hua Wang, Shalini Chopra, Nima Hooshdaran, Hyunjung Kim, Jie Zhuo, Shion A. Lim, Jianing Yang, Kevin K. Leung, Emily Egusa, Li Zhang, Adam Foye, Jonathon Chou, Felix Y. Feng, Eric J. Small, James A. Wells, Rahul Aggarwal, Michael J. Evans

## Abstract

**Purpose:** Radioligand therapy (RLT) is relatively unexplored in metastatic castration resistant prostate cancer (mCRPC), with much of the focus having been on bone seeking radionuclides and PSMA-directed RLT. Herein, we evaluated if CUB domain containing protein 1 (CDCP1) can be exploited to treat mCRPC with RLT, particularly for subsets like small cell neuroendocrine prostate cancer (SCNC) that would not be expected to respond to current options.

**Experimental Design:** *CDCP1* mRNA levels were evaluated in the RNA-seq data from 119 recent mCRPC biopsies. Protein expression was assessed in twelve SCNC and adenocarcinoma patient derived xenografts. Saturation binding assays were performed with 4A06, a recombinant human antibody that targets the CDCP1 ectodomain. The feasibility of imaging and treating mCRPC in vivo was tested with ^89^Zr-4A06 and ^177^Lu-4A06.

**Results:** *CDCP1* mRNA expression was observed in over 90% of mCRPC biopsies, including SCNC and in adenocarcinoma with low *FOLH1* (PSMA) levels. A modest anticorrelation was observed between *CDCP1* and *PTEN*. Overall survival was not significantly different based on *CDCP1* mRNA levels, regardless of PTEN status. Full length and/or cleaved CDCP1 was expressed in ten of twelve PDX samples. Bmax values of ~22,000 and ~6,200 fmol/mg were calculated for two human prostate cancer cell lines. Five prostate cancer models were readily detected in vivo with ^89^Zr-4A06. ^177^Lu-4A06 significantly suppressed the growth of DU145 tumors compared to control.

**Conclusions:** The antitumor data and the overexpression of CDCP1 reported herein provide the first evidence promoting CDCP1 directed RLT as a treatment strategy for mCRPC.

**Statement of Translational Relevance:** New targets for RLT are needed to address the subset of mCRPC that cannot be treated with bone seeking radionuclides or PSMA directed RLT. We report herein the first data credentialing CDCP1 as a target for mCRPC, in both adenocarcinoma and neuroendocrine subtypes. Combined with low expression in normal human tissues, these data provide a compelling scientific rationale for testing CDCP1 directed RLT clinically in mCRPC patients alone or in combination with other systemic therapies.

## Introduction

RLT is currently undergoing a renaissance, fueled in part by advances in “-omic” technologies to better identify suitable targets, a stronger understanding of the features of ligand design required to minimize dose limiting toxicity, and a larger repertoire and supply of therapeutically beneficial radioisotopes, including α particle emitters (e.g. Ac-225, Ra-223). Moreover, the recent FDA approvals of Lutathera and Azedra, which target unrelated proteins in very different tumor types, combined with encouraging clinical activity for experimental RLTs engaging diverse cancer antigens (e.g. PSMA, HER2, CXCR4) and malignancies, show a versatility to RLT that suggests this modality may be a scalable approach for the treatment of many deadly cancers(1–3).

RLT as a treatment approach for mCRPC is still relatively unexplored, with much of the community’s attention having been paid to PSMA-directed RLT and bone seeking radionuclides. Even with this small sample size, the clinical data suggest that RLT can be a viable treatment option for mCRPC. For example, ^223^RaCl_2_ (aka Xofigo) was granted FDA approval based on the ALSYMPCA trial, and ^177^Lu-PSMA 617 has demonstrated promising activity in multiple prospective studies, with rapid tumor responses observed in the majority of patients, including those with treatment-refractory disease (4,5). That said, these RLTs have shortcomings that limit their impact on mCRPC treatment. Xofigo obviously cannot treat soft tissue tumors, and responses in bony metastases are transient. Tumor response to PSMA-directed RLT is also limited in duration, in part due to dose limiting toxicity (e.g. cytopenias, xerostomia), intrinsic or acquired tumor resistance, and the heterogeneity of PSMA expression in mCRPC (6). PSMA-directed RLT is also not suitable for treating low-PSMA expressing mCRPC including small cell neuroendocrine cancer (SCNC), which is an increasingly common subset of mCRPC observed in 15-20% of cases (7). Collectively, these observations provide a strong scientific rationale for exploring new RLTs to improve treatment outcomes in mCRPC.

We hypothesized that the single pass transmembrane protein CUB domain containing protein 1 (CDCP1) could be potential target for RLT that overcomes some of the abovementioned limitations. For example, after two reports between 2008-2015 showed CDCP1 overexpression in human prostate cancer cell lines and small cohorts of primary tumors, a recent study from Alimonti et al. has more convincingly established that CDCP1 overexpression is a common feature of prostate cancer (8–10). Mining open access RNA-seq data from tumor biopsies, as well as determining protein expression in biopsies with immunohistochemistry showed that CDCP1 is overexpressed in ~50% of mCRPC biopsies and ~30% of primary tumors. Interestingly, overexpression co-occurred in tumors with PTEN loss and was observed in tumors with a more aggressive phenotype. Further supporting CDCP1 as a target for RLT, its expression is generally low in normal human tissues, with the highest levels observed in breast tissue (11,12). This restricted expression distinguishes CDCP1 from PSMA which is broadly expressed in the salivary gland, liver, kidney, and other tissues. Lastly, we recently established proof of concept for CDCP1 directed RLT by showing that that pancreatic cancer tumors can be effectively treated with an anti-CDCP1 antibody termed 4A06 coupled to β- or α emitters (13). On this basis, we endeavored herein to explore the feasibility of treating mCRPC with CDCP1-directed RLT.

## Material and Methods

### General Methods

All materials and chemicals were purchased from commercial vendors and used without further processing/purification. DU145, 22RV1 and PC3 Cell lines were procured from American Type Tissue Collection (ATCC) and cultured according to manufacturer’s recommendations. The monoclonal antibody 4A06 was expressed and purified in the IgG1 format as previously described (12). p-SCN-Bn-Deferoxamine (B-705) and p-SCN-Bn-DOTA (B-205) were purchased from Macrocyclics (Plano, TX) and directly used for conjugation without further purification to antibody. ^89^Zr-oxalate was obtained from 3D Imaging, LLC (Maumelle, AR). ^177^LuCl_3_ was obtained from Oak Ridge National Laboratory. Iodine-125 was obtained from Perkin Elmer.

### Analysis of RNA-seq data

A recent biopsy gene expression dataset from mCRPC patients clinically annotated with survival data was used for analysis (14,15). Patient samples were obtained using image-guided core needle biopsy of metastatic lesion in the bone or soft tissue, with cores freshly frozen for RNA sequencing. SCNC status was determined via unsupervised hierarchical clustering using a previously validated gene signature (15). The following list of genes was used for hierarchical clustering to determine SCNC status: AR, TMPRSS2, GATA2, HOXB13, KLK3, FOXA1, NKX3-1, CHGB, FOXA2, SOX2, SCG2, NKX2-1, REST, SPDEF, NOTCH2, NOTCH2NL, ASCL1, ETV1, ETV4, ETV5, RB1, CDKN2A, E2F1. Linear regression between *CDCP1* and *FOLH1* expression was performed in R (v4.0.2), with the resulting P value and Pearson correlation coefficient reported.

### Flow Cytometry

Actively proliferating cells were lifted mechanically from a tissue culture plate and placed in the primary antibody solution diluted in PBS (4A06, 1:1,000). After a 30 min incubation, the cells were washed with PBS multiple times and placed in a fluorophore conjugated secondary antibody solution for 30 minutes on ice (1:500, 109-546-097, Jackson ImmunoResearch). Cells were collected and washed before being placed in PBS and passed through a cell strainer. Samples were taken to a flow machine (FACS CantoII). Data analysis was performed using FloJo.

### Immunoblot

Tumor samples were added to Pierce RIPA lysis buffer (89900, ThermoFisher Scientific) with Halt protease and phosphatase inhibitors (1861281, ThermoFisher Scientific). The samples were homogenized using a probe sonicator (Omni TH-01) and then centrifuged for 15 minutes at 15,000 × g. 4X SDS-loading buffer was added to protein lysates and 15 μg of lysate were resolved on a Bolt 4-12% gel (NW04120BOX, Invitrogen). Gels were transferred onto an Immobilon-P membrane (IPVH00010, Millipore). Membranes were blocked in 5% milk in TBST for 90 minutes before being placed in a primary antibody solution. Primary antibodies used were CDCP1 (4115S, Cell Signaling, 1:1000), PSMA (12815S, Cell Signaling, 1:1000), AR (515S3, Cell Signaling, 1:1000), and B-Actin (A5441, Sigma-Aldrich, 1:5000). Membranes were then washed and incubated with a secondary antibody solution for 30 min at room temperature. Secondary antibodies used were goat α-rabbit (65-6120, Invitrogen, 1:5000) and goat α-rat (62-6520, Invitrogen, 1:5000). Proteins were detected using West Pico Chemiluminescent Substrate for 20 seconds (34578, ThermoFisher Scientific) and then exposed to film (30-507, Blue Devil).

### Saturation binding assays

To prepare ^125^I-4A06, 6.0 μL of ^125^I in 1M NaOH (~ 340 uCi) was added into a vial containing 100.0 μL of HEPES buffer (0.5 M). This solution was transferred to an iodination tube (Peirce) and 4A06 antibody (300μg) was then added. The tube was incubated for 10 min at room temperature, and ITLC showed 85.0% radiolabeling efficiency (solvent: 20nM citric acid). ^125^I-4A06 was purified using a G-25 column. The final yield of the purified ^125^I-4A06 was 68.58% (specific activity =1.13 μCi/μg) and purity was > 99%.

DU145 and PC3 (0.6 × 10^6^ cell/well) were seeded on 12-well plates using DMEM (10% FBS). The cells were harvested, washed, and suspended in PBS for the saturation binding assay. Total binding of ^125^I-4A06 was determined by adding it to cell suspensions at seven concentrations from 0.025 nM to 10 nM. Non-specific binding was determined by adding 1000x cold 4A06 to ^125^I-4A06/cell mixtures at 0.025nM, 0.3 nM, and 10 nM. In all cases, cells were incubated with ^125^I-4A06 at room temperature for 1 hr, washed with PBS, and lysed by adding 1.0 M NaOH. The bound and unbound radioactive fractions were collected and measured on a Hidex Gamma counter (Turku, FI) to determine Bmax.

### Animal studies

All animal studies were conducted in compliance with Institutional Animal Care and Use Committee at UCSF. For tumor imaging or treatment studies with mCRPC xenografts from cell line implants, four to six-week-old intact male athymic nu/nu mice (Charles River) were utilized. Animal studies involving PDX’s utilized four to six week old intact NOD SCID gamma mice (Charles River). For tumor implantation, mice were inoculated subcutaneously (~1.5 ×10^6^ cells) in the flank. The cells were injected in a 1:1 mixture (v/v) of media (DMEM) and Matrigel (Corning). Xenografts were generally palpable within 3-4 weeks after injection. Mice bearing subcutaneous DU145 tumors received ^177^Lu-4A06 (400 μCi) or vehicle (saline) via tail vein at day 0 and day 5 of the study period. Animals were weighed at the time of injection, and once weekly until the completion of the study. Tumor volume measurements were calculated with calipers. The study endpoints were death due to tumor volume >1000 mm^3^ or ≥ 20% loss in mouse body weight.

### Small animal PET/CT

All data, including the raw imaging files, are available upon request. 4A06 was functionalized with desferrioxamine (DFO) and subsequently radiolabeled with Zr-89 as previously described (13). Tumor-bearing mice received ~200 μCi of ^89^Zr-4A06 in 100 μL saline solution volume intravenously using a custom mouse tail vein catheter with a 28-gauge needle and a 100-150 mm long polyethylene microtubing. Mice were imaged on a small animal PET/CT scanner (Inveon, Siemens Healthcare, Malvern, PA). Animals were typically scanned for 30 minutes for PET, and the CT acquisition was performed for 10 minutes.

The co-registration between PET and CT images was obtained using the rigid transformation matrix generated prior to the imaging data acquisition since the geometry between PET and CT remained constant for each of PET/CT scans using the combined PET/CT scanner. The photon attenuation was corrected for PET reconstruction using the co-registered CT-based attenuation map to ensure the quantitative accuracy of the reconstructed PET data. Decay corrected images were analyzed using AMIDE software.

### Small animal SPECT/CT

4A06 was functionalized with DOTA and radiolabeled with Lu-177 as previously described (13). Mice received an intravenous injection of ^177^Lu-4A06 (~400 μCi/mouse) at day 0 and day 5 of the antitumor assessment study. At 48 hours post injection of the second dose, the mice were imaged under anesthesia using a small animal SPECT/CT (VECTor4CT, MILabs, Utrecht, The Netherlands). Animals were typically scanned for 40 minutes for SPECT, and the CT acquisition was performed for 10 minutes. The co-registered CT was used for photon attenuation correction in the SPECT reconstruction. The decay corrected images were reconstructed and analyzed using AMIDE software.

### Biodistribution studies

At a dedicated time after radiotracer injection, animals were euthanized by cervical dislocation. Blood was harvested via cardiac puncture. Tissues were removed, weighed and counted on a Hidex automatic gamma counter (Turku, Finland). The mass of the injected radiotracer was measured and used to determine the total number of counts per minute by comparison with a standard of known activity. The data were background- and decay-corrected and expressed as the percentage of the injected dose/weight of the biospecimen in grams (%ID/g).

### Digital autoradiography

Post mortem, tumors were harvested, transferred to sample boats, and immersed in Tissue–Plus OCT compound (Scigen, Gardena, CA). The tissues were snap frozen at −80° C. The tumor tissue was sectioned into 20 μm slices using a microtome (Leica, Buffalo Grove, IL) and mounted on glass microscope slides. The slides were loaded onto an autoradiography cassette and exposed with a GE phosphor storage screen for 24-72 hours at −20° C. The film was developed and read on a Typhoon 9400 phosphorimager (Marlborough, MA). <u>Images</u> of whole sections were acquired on a VERSA automated slide scanner (Leica Biosystems, Wetzlar, Germany), equipped with an Andor Zyla 5.5 sCMOS camera (Andor Technologies, Belfast, UK). ImageScope software (Aperio Technologies, Vista, CA) is used for creating individual images. Photoshop CS6 software (Adobe Systems, McLean, VA) were used for montage and processing. The autoradiography and histology were analyzed using ImageJ.

### Statistical analysis

Data were analyzed using PRISM v8.0 software (Graphpad, San Diego, USA). Binary comparisons between two treatment arms were made with an unpaired, two-tailed Student’s t-test. Differences at the 95% confidence level (P < 0.05) were considered to be statistically significant. Analysis of statistically significant changes in Kaplan Meier survival curves were made with a log rank test. Differences at the 95% confidence level were considered statistically significant. Sample size was estimated based on an expected effect size of 0.2 for tumor volume changes with a type I error rate of 0.05.

Overall survival was measured from the date of metastatic biopsy in the mCRPC patients. Kaplan-Meier product limit method and log-rank test were used to investigate the relationship between *CDCP1* expression (transcripts per million, TPM) and overall survival, or *CDCP1* expression, *PTEN* mutation status and overall survival. The patient cohort was dichotomized above and below median *CDCP1* expression as well as broken into quartiles of *CDCP1* expression for survival analyses.

## Results

### CDCP1 is expressed in mCRPC, including SCNC and PSMA low expressing adenocarcinoma

Following on the expression analysis from Alimonti et al, we probed our recently published mCRPC data set to assess *CDCP1* expression (15). *CDCP1* was expressed in 90% of mCRPC biopsies (107 of 119 samples, see **Figure 1A**). Since previous analyses showed that CDCP1 mRNA and protein levels are upregulated in PTEN null prostate cancer, we tested for a correlation between *CDCP1* and *PTEN* mRNA. A Pearson analysis revealed that *CDCP1* levels were modestly, but significantly, anti-correlated (**Figure 1B**). We also compared *CDCP1* mRNA levels between patients with wild type PTEN and patients with any type of PTEN mutation (i.e. deletion, inversion, break-end). We found that *CDCP1* is slightly upregulated in PTEN-mutant patients, though the effect is not statistically significant (**Table 1**).

**Figure 1.**
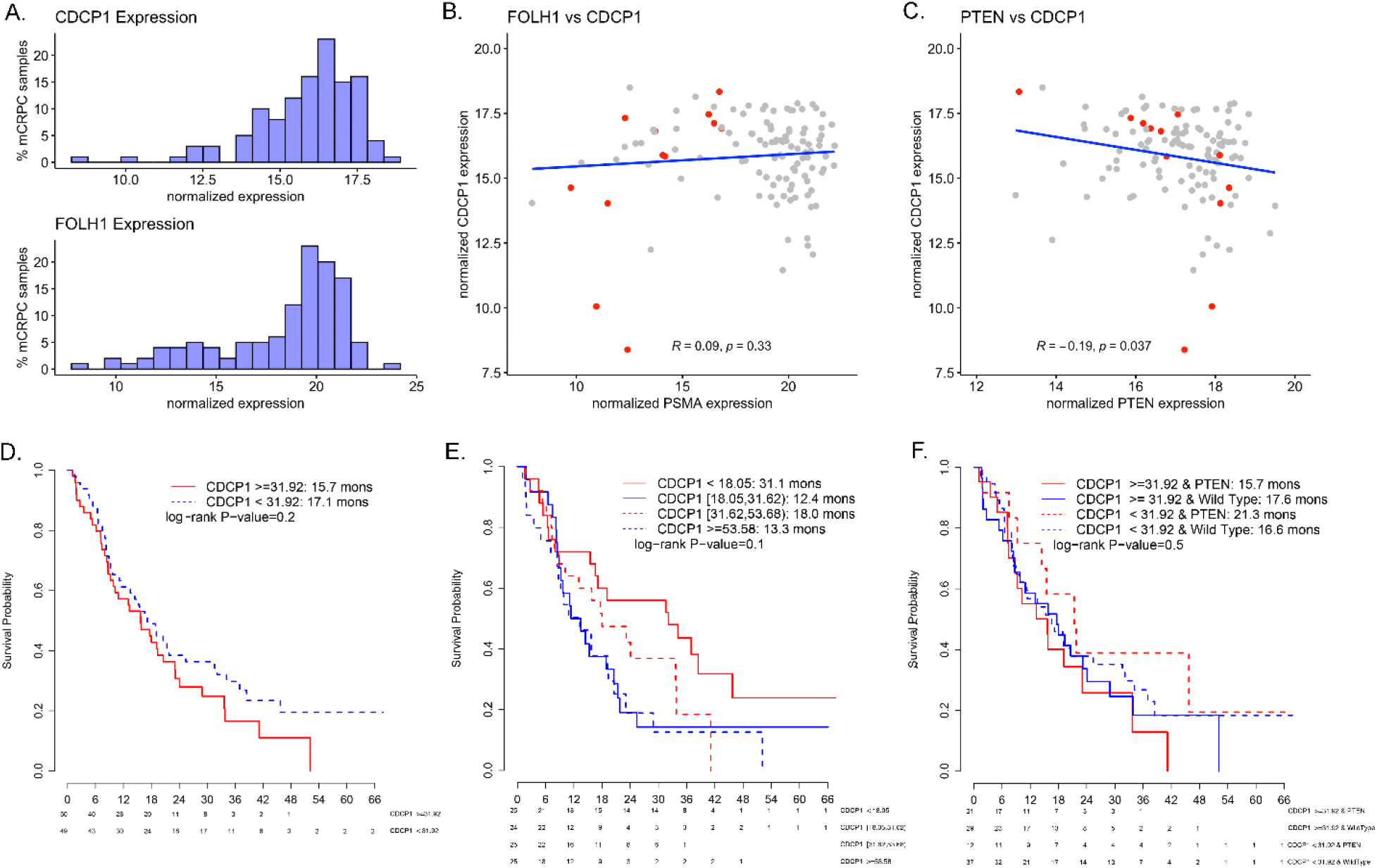
An expression analysis of CDCP1 from RNA-seq data collected in mCRPC biopsies. **A.** Histograms showing the distribution of *CDCP1* mRNA levels over the set of 114 mCRPC biopsies. *FOLH1* (PSMA), an abundantly expressed target for RLT, is shown for comparison. **B.** A scatter plot showing the expression of *CDPC1* and *FOLH1* per biopsy. A Pearson correlation shows no significant trend in expression between the two genes. NEPC tumors are highlighted in red. **C.** A scatter plot showing the expression of *CDPC1* and *PTEN* per biopsy. A Pearson correlation shows the genes are significantly anti-correlated, consistent with previous analyses of primary and metastatic prostate cancer biopsies. SCNC tumors are highlighted in red. **D.** Kaplan Meier curves showing the average survival in dichotomized cohorts above and below median *CDCP1* expression. Despite previous reports indicating that *CDCP1* overexpression correlates with shorter overall survival, no significant difference was observed in this data set. **E.** Kaplan Meier curves showing the average survival in cohorts broken into quartiles of *CDCP1* expression. No significant difference in overall survival was observed between the groups. F. Kaplan Meier curves showing the average survival in cohorts grouped by *CDCP1* expression (above and below median) and *PTEN* status (wild type versus mutant). No significant difference was observed between groups.

**Table 1.** A summary of the results from a differential gene expression analysis to compare expression of CDCP1 between patients with wild type PTEN and patients with any type of PTEN mutation (deletion, inversion, break-end). CDCP1 is slightly upregulated in PTEN-mutant patients, but this is not statistically significant.

| Gene | Log2 Fold Change | Unadjusted P value | False discovery rate |
| --- | --- | --- | --- |
| PTEN | -1.44 | $2.1 \times 10^{-6}$ | 0.04 |
| CDCP1 | 0.52 | 0.20 | 0.88 |

We next evaluated the distribution of *CDCP1* versus *FOLH1* (PSMA) expressing tumors and found that 93 of 119 (78%) were predicted to be mutually positive, while 3 of 119 (3%) were mutually negative. The data set also showed that 14 of 119 (12%) were predicted to have *CDCP1* expression but not *FOLH1* expression, while 9 of 119 (8%) were predicted to be *FOLH1* positive but *CDCP1* negative. We compared the expression of *CDCP1* and *FOLH1* expression per patient and found that no significant correlation based on a Pearson analysis (**Figure 1C**).

We found no relationship between *CDCP1* expression and overall survival in dichotomized cohorts above and below median expression (log-rank p value = 0.2; **Figure 1D**) or by breaking into quartiles of *CDCP1* expression (log-rank p-value = 0.1; **Figure 1E**). When subdividing the patient cohort into subgroups based upon *CDCP1* expression (above vs. below median) and *PTEN* mutation status (wild type vs. mutation), there was also no difference in survival between subgroups (log-rank p-value = 0.5; **Figure 1F).**

Since CDCP1 protein was already found to be expressed at high levels in the whole cell lysates of several human prostate cancer cell lines (10), we evaluated expression in human PDX whole cell lysates. Full length and/or cleaved CDCP1 was expressed in four of five adenocarcinoma PDXes that we evaluate from the LuCaP and Living Tumor Laboratory series (**Figure 2A**). The highest levels were observed in LuCaP70 CR and LTL-484, an example of adenocarcinoma with negligible PSMA expression. Remarkably, expression was detected in six of seven SCNC PDXs, with the highest expression observed in LTL-545 and LTL-370 (**Figure 2B**). In line with some recent reports suggesting that proteolytically cleaved CDCP1 may contribute to early tumor development and metastatic potential (16–18), CDCP1 was detected as both the full length and cleaved forms.

**Figure 2.**
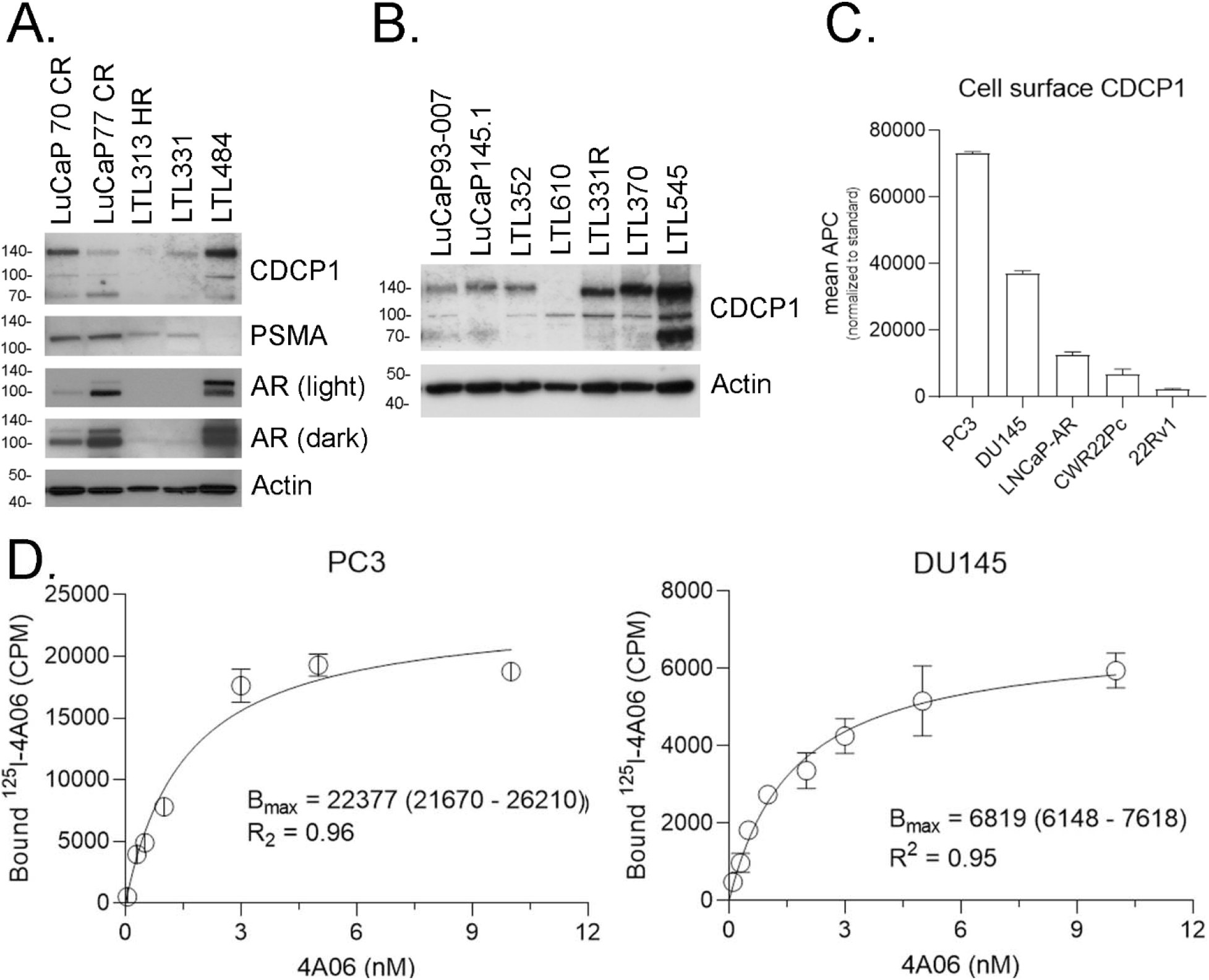
CDCP1 is abundantly expressed in human prostate cancer models, including PSMA low adenocarcinoma and SCNC. **A.** Immunoblot data showing CDCP1 expression in human prostate cancer adenocarcinoma PDX whole cell lysates. **B.** Immunoblot data showing CDCP1 expression in human SCNC prostate cancer PDX whole cell lysates. **C.** Flow cytometry data showing the relative levels of total, cell surface CDCP1 (full length and cleaved) in five human prostate cancer cell lines. D. Saturation binding assays with ^125^I-4A06 performed on PC3 and DU145 cells. The B_max_ (95% confidence intervals) values are reported inset for each study.

We next confirmed CDCP1 expression on the surface of viable prostate cancer cells with flow cytometry. The relative cell surface levels in five human prostate cancer cell lines (PC3 > DU145 > LNCaP-AR > CWR22Pc > 22Rv1) aligned with previously reported total expression levels on immunoblot (**Figure 2C** and **Supplemental Figure 1**)(10). To more rigorously quantify CDCP1 abundance in mCRPC, we performed saturation binding assays to quantify receptor number per cell using 4A06, a monoclonal recombinant human antibody that recognizes a region of the CDCP1 ectodomain present in both full length and cleaved CDCP1. We calculated PC3 cells to have a B_max_ of ~22,000 fmol/mg (~1 × 10^7^ receptors per cell) and DU145 to have a B_max_ of ~6500 fmol/mg (~1 × 10^6^ receptors per cell, see **Figure 2D**). These values exceed receptor densities reported for PSMA in human prostate cancer cell lines (~1 × 10^5^ receptors/cell) (19).

### Tumor autonomous expression of CDCP1 in mCRPC is detectable with ^89^Zr-4A06

We next evaluated if CDCP1 expression can be detected on PET using ^89^Zr-labeled 4A06. 4A06 was functionalized with desferrioxamine B and radiolabeled to high yield with Zr-89 as previously reported (13). A cohort of mice (n = 4/tumor type) bearing one of the following tumors were studied: TRAMP C2, DU145, 22Rv1, LTL-545 and LTL-331. The tumors represent a mixture of adenocarcinoma (TRAMP C2, 22Rv1, LTL-331) and androgen receptor null mCRPC or NEPC (DU145, LTL-545). ^89^Zr-4A06 PET conducted 48 hours post injection showed clear evidence of radiotracer accumulation in tumors (**Figure 3A**). Post mortem biodistribution measurements at 48 hours post injection showed that TRAMP C2 and DU145 had the highest radiotracer uptake at 16.65 ± 1.6 and 12.9 ± 1.9% ID/g, respectively (**Figure 3B** and **Supplemental Figure 2**). The tumor to muscle ratios were at least 7:1 and DU145 had the highest ratio at ~25:1 (**Figure 3C**). Lastly, we performed digital autoradiography to assess radiotracer distribution within a tumor representing high (DU145) and comparatively lower (22Rv1) tracer uptake. We found that radiotracer distribution was qualitatively similar, and most abundant around the periphery of the tumor as expected for a large immunoglobulin like 4A06 (**Figure 3D**).

**Figure 3.**
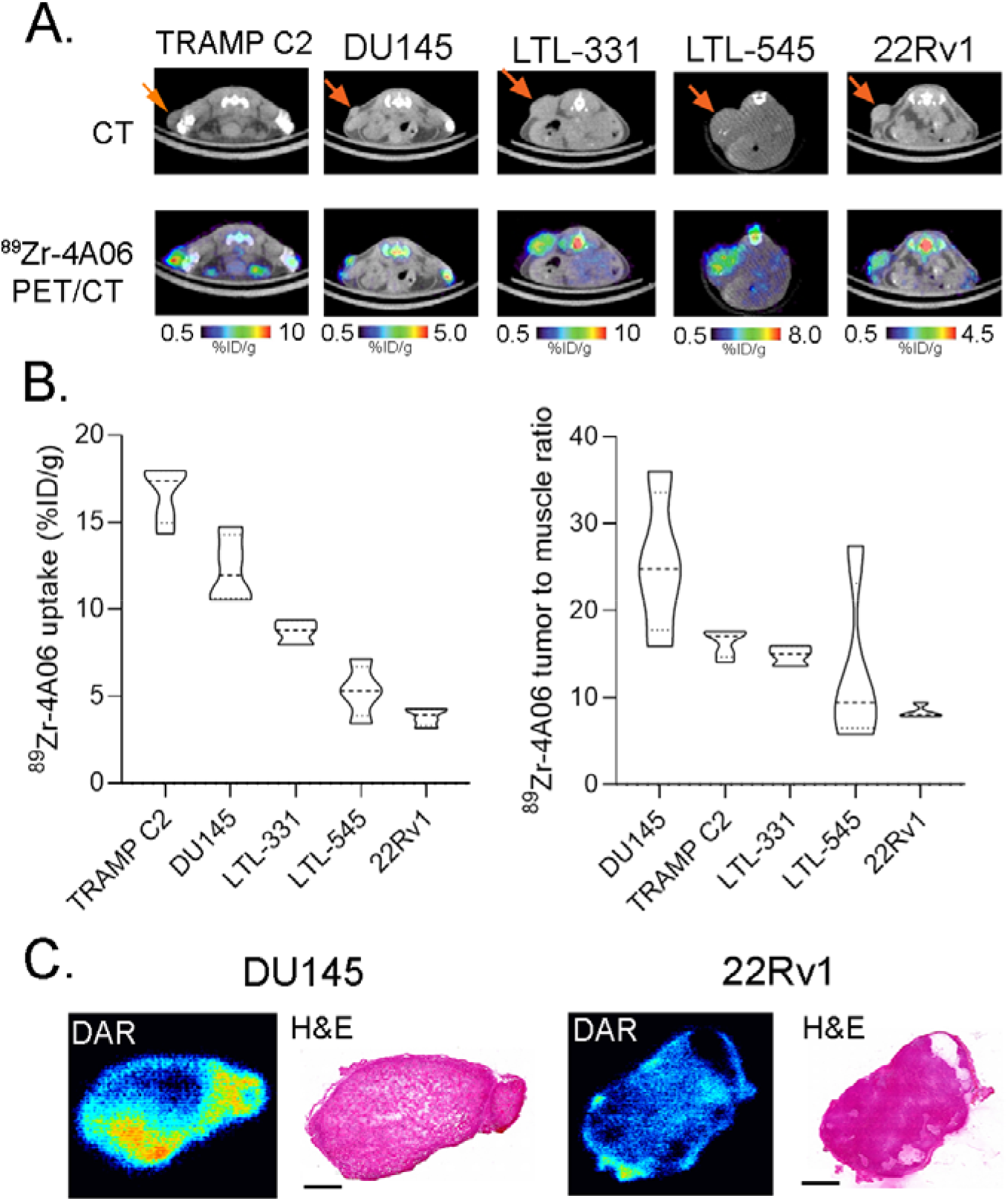
Tumor autonomous CDCP1 expression is measurable in vivo with ^89^Zr-4A06 PET. **A.** Transaxial CT and PET/CT images showing the tumoral uptake of ^89^Zr-4A06 at 48 hours post injection. The arrows indicate the position of the subcutaneous tumor. **B.** Biodistribution data showing the tumoral uptake of ^89^Zr-4A06 at 48 hours post injection. **C.** Tumor to muscle ratios collected at 48 hours post injection. **D.** Autoradiography showing the distribution of ^89^Zr-4A06 within DU145 and 22Rv1 tissue slices.

### Radioligand therapy with ^177^Lu-4A06 inhibits mCRPC tumor growth

We next tested if CDPC1 can be targeted for RLT in mCRPC tumors. Mice bearing PSMA-negative DU145 tumors received ^177^Lu-4A06 in two fractions of 400 μCi at day 0 and day 5 of the study. We chose this dosing regimen as we previously showed it to be more efficacious that a single fraction of an equivalent total radioactive dose (i.e. 800 μCi) (13). SPECT/CT imaging showed effective tumor targeting at 72 hours after the first injection (**Figure 4A**). Following the tumor volume changes for 60 days showed that ^177^Lu-4A06 significantly inhibited tumor growth compared to vehicle (*P* < 0.01, **Figure 4B** and **Supplemental Figure 3**). Moreover, ^177^Lu-4A06 was well tolerated, as expected (**Supplemental Figure 4**).

**Figure 4.**
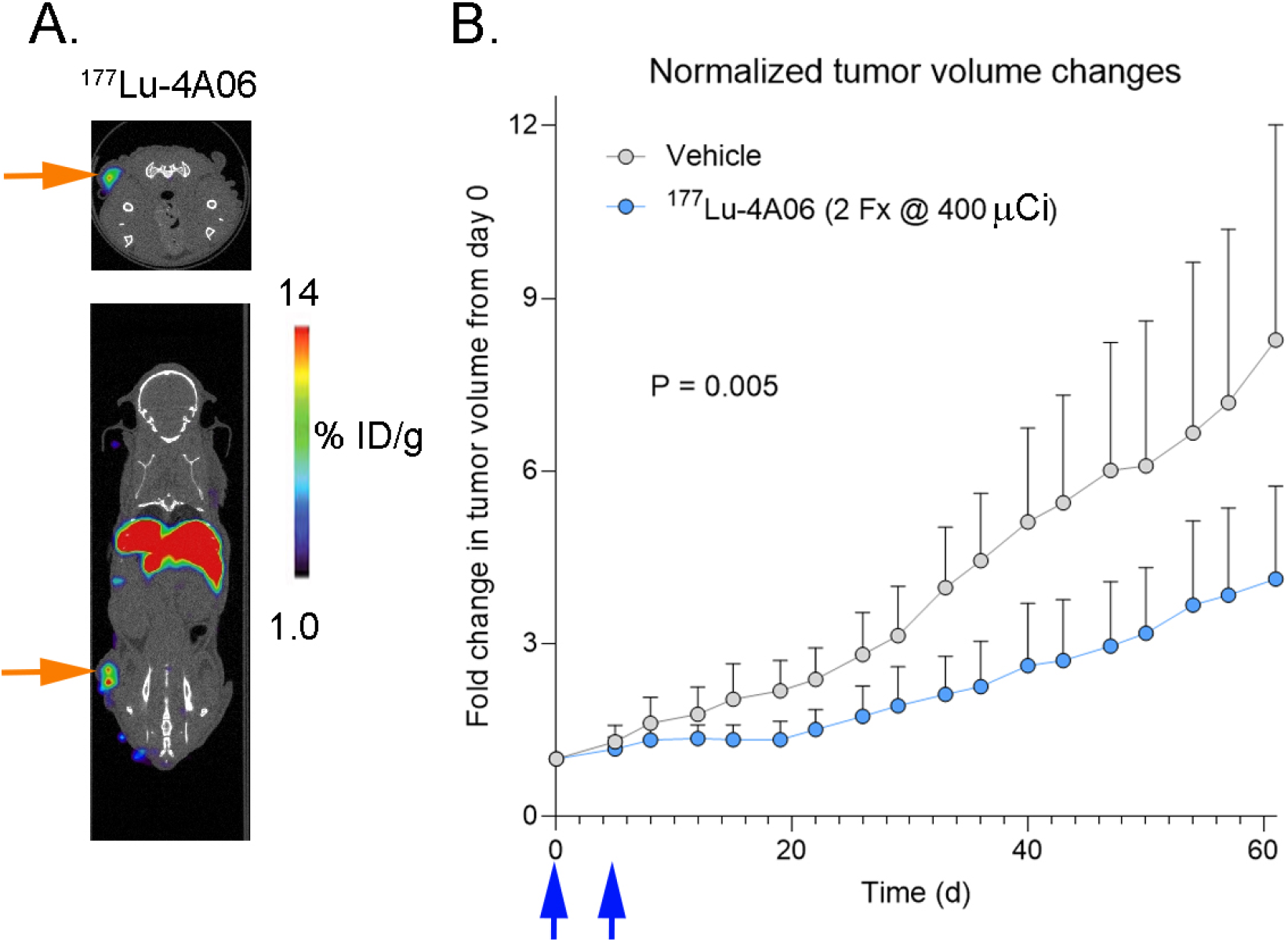
^177^Lu-4A06 inhibits the growth of a PSMA null prostate cancer tumor. **A.** A SPECT/CT image showing tumoral localization of ^177^Lu-4A06 in a mouse bearing a subcutaneous DU145 tumor (orange arrow). The image was acquired at 72 hours after a second injection of 400 μCi of ^177^Lu-4A06 (i.e. 10 days after the start of the study). **B.** Tumor volume data showing the suppression of tumor growth by ^177^Lu-4A06, administered in two fractions on day 0 and day 7 (blue arrows). The difference in tumor growth between treatment arms (n = 8/arm) was significant, P<0.01.

## Discussion

In this report, we demonstrate that CDCP1 directed RLT may be a potential therapeutic strategy for the treatment of mCRPC. We found that CDCP1 expression was observed in 90% of mCRPC biopsies, including SCNC and in adenocarcinoma with negligible PSMA expression. Moreover, CDCP1 was anticorrelated with PTEN expression, consistent with prior analyses of separate prostate cancer biopsy data sets. Full length and/or cleaved CDCP1 was expressed in six of seven SCNC PDX samples and expressed in four of five adenocarcinoma PDX samples, including LTL-484 which has low PSMA expression. The feasibility of detecting tumor autonomous CDCP1 expression was demonstrated by applying ^89^Zr-4A06 PET to five subcutaneous prostate cancer mouse models. Lastly, an antitumor assessment study showed that ^177^Lu-4A06 significantly suppressed the growth of PSMA null DU145 tumors compared to control and extended overall survival.

The broad expression of CDCP1 in both adenocarcinoma and SCNC raises the question as to what signaling pathway(s) promote CDCP1 overexpression. One possibility is that CDCP1 expression may be elevated transcriptionally by one of the hypoxia inducible factors, as CDCP1 was previously found to be a HIF2α target gene and HIFs can be activated downstream of mTOR (CDCP1 overexpression appears to associate with PTEN null tumors) (20–22). Mechanism studies are currently underway, which may also lead to rational combination treatment strategies with CDCP1 directed therapeutics.

A role for CDCP1 expression in prostate cancer has yet to be fully defined. Early functional data suggest CDCP1 may have oncogenic properties, especially in the context of PTEN loss, but the molecular details underpinning this observation require further elaboration (8). Our immunoblot data, combined with previous analyses in cell line models, do not appear to suggest that prostate cancer preferentially expresses the full length versus cleaved form of CDCP1. Thus, CDCP1 may have several functions in prostate cancer pathobiology, as tumor signaling functions unique to the cleaved form of CDCP1 have been recently discovered. Deeper analysis of CDCP1 proteolytic isoforms in patient biopsies is required to define the overall incidence of cleaved versus full length CDCP1.

mCRPC is biologically heterogeneous, which increases the urgency to expand the current repertoire of RLTs and their corresponding targets to maximize disease coverage. In this regard, our preliminary findings that CDCP1 is expressed in PSMA null mCRPC provides a compelling scientific rationale to investigate expression at the protein level in patient biopsies, as well as translate an anti-CDCP1 radioligand to determine patient dosimetry. We are actively working on both frontiers.

Lastly, these data come at an exciting time, as several classes of therapeutics that logically synergize with ionizing radiation have achieved regulatory approval for prostate cancer treatment. Examples include the PARP inhibitor olaparib and antiandrogens like enzalutamide and apalutamide that repress DNA damage repair machinery but also may induce CDCP1 expression. The nuclear medicine community is just beginning to explore in patients the feasibility and effiicacy of combination treatments with RLTs, and the early antitumor data are very encouraging.(23–26) Thus, combining CDCP1 directed RLT with standard of care treatments for mCRPC may be a potential strategy to achieve deeper clinical responses.

## Author Contributions

**Conception and design:** N. Zhao, L. Zhang, J. Chou, J.A. Wells, R. Aggarwal, M.J. Evans

**Acquisition of data:** N. Zhao, K. Trepka, Y. Huang, S. Chopra, N. Hooshdaran, H. Kim, L. Zhang, J. Zhuo, J. Yang, S.A. Lim, J. Zhuo, A. Foye, KK Leung

**Analysis and interpretation of data:** N Zhao, L. Zhang, J. Chou, F.Y. Feng, E.J. Small, J.A. Wells, R. Aggarwal, M.J. Evans

**Writing, review, and/or revision of the manuscript:** N. Zhao, R. Aggarwal, M.J. Evans

## Acknowledgments

We gratefully acknowledge Dr. Henry VanBrocklin, Dr. Denis Beckford-Vera, and Joseph Blecha for assistance with radiochemistry and Dr. Youngho Seo and Ryan Tang for assistance with small animal PET/CT studies. Small animal PET/CT and SPECT/CT studies were performed on instruments supported by National Institutes of Health grants S10RR023051 and S10OD012301. This research was supported by a Stand Up To Cancer-Prostate Cancer Foundation Prostate Cancer Dream Team Award, grant number SU2C-AACR-DT0812 (PI: EJS). Stand Up To Cancer is a division of the Entertainment Industry Foundation. This research grant was administered by the American Association for Cancer Research, the scientific partner of SU2C. J.A.W. acknowledges funding from NCI 1P41CA196276, CA191018, NIH GM097316; and Celgene Corp. S.L. was supported by Helen Hay Whitney post-doctoral fellowship. J.Z. was supported by an NIH post-doctoral fellowship. M. J. Evans acknowledges funding from the American Cancer Society (Research Scholar Grant 130635-RSG-17-005-01-CCE), the Precision Imaging of Cancer and Therapy program at UCSF, the Department of Defense (W81XWH-18-1-0763) and the National Institutes of Health (R01MH114053, R01EB025027).

