## Supplemental Material for "CUB domain containing protein 1 (CDPC1) is a target for radioligand therapy in castration resistant prostate cancer"

**Supplemental Figure 1.** Representative histograms showing the relative CDPC1 expression levels on the surface of human prostate cancer cell lines. The experiments were performed in triplicate to arrive at the data presented in Figure 2B. In culture, CWR22Pc is an inseparable mixture of human prostate cancer cells and mouse fibroblasts which is reflected in the bimodal distribution of the histogram.

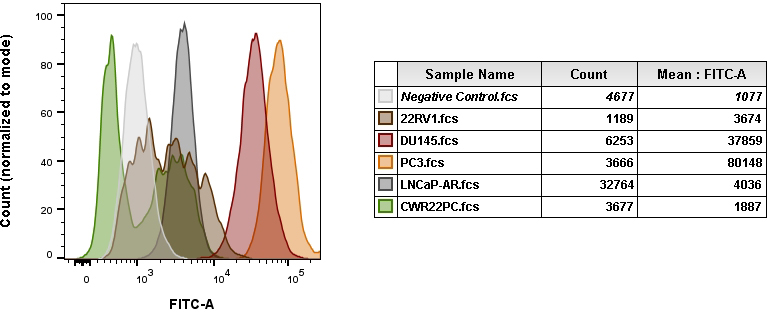

**Supplemental Figure 2.** A summary of the biodistribution data collected from tumor and selected normal organs at 48 hours post injection of ^89^Zr-4A06. Male nu/nu mice were the host for DU145, 22Rv1, and PC3, while NSG mice were used for LTL-545 and LTL-331.

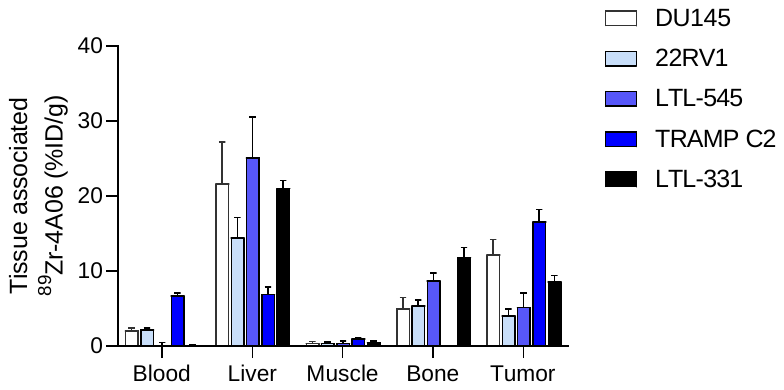

**Supplemental Figure 3.** Individual tumor volumes from the antitumor assessment study in nu/nu male mice bearing DU145 xenografts and treated with vehicle or ^177^Lu-4A06 (2 Fx @ 400 μCi/Fx over 8 days).

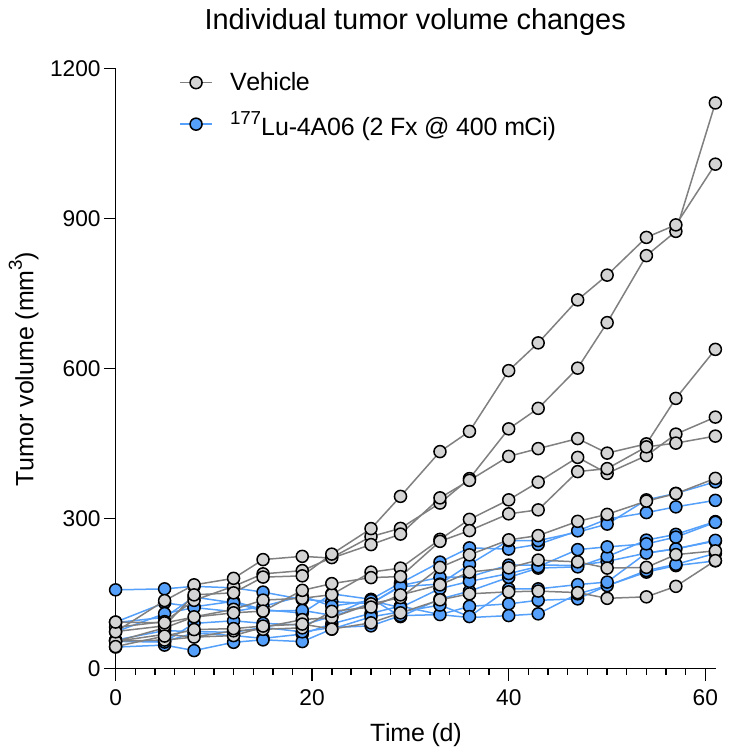

**Supplemental Figure 4.** Body weights from the male nu/nu mice subjected to the antitumor assessment study with ^177^Lu-4A06 or vehicle.

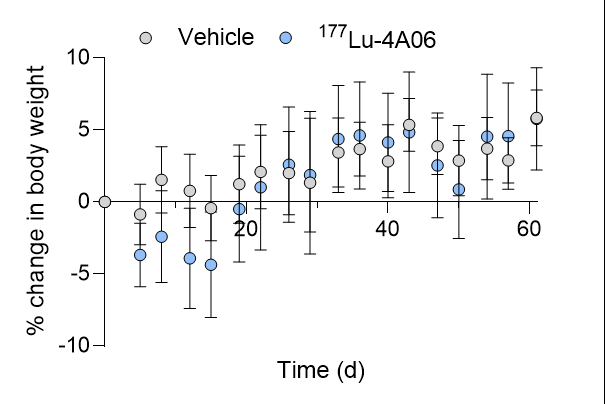
